# Exploring the metabolic landscape of pancreatic ductal adenocarcinoma cells using genome-scale metabolic modeling

**DOI:** 10.1101/2021.07.14.452356

**Authors:** Mohammad Mazharul Islam, Andrea Goertzen, Pankaj K. Singh, Rajib Saha

## Abstract

Pancreatic ductal adenocarcinoma (PDAC) is a major research focus due to its poor therapy response and dismal prognosis. PDAC cells adapt their metabolism efficiently to the environment to which they are exposed, often relying on diverse fuel sources depending on availability. Since traditional experimental techniques appear exhaustive in the search for a viable therapeutic strategy against PDAC, in this study, a highly curated and omics-informed genome-scale metabolic model of PDAC was reconstructed using patient-specific transcriptomic data. From the analysis of the model-predicted metabolic changes, several new metabolic functions were explored as potential therapeutic targets against PDAC in addition to the already known metabolic hallmarks of pancreatic cancer. Significant downregulation in the peroxisomal fatty acid beta oxidation pathway reactions, flux modulation in the carnitine shuttle system, and upregulation in the reactive oxygen species detoxification pathway reactions were observed. These unique metabolic traits of PDAC were then correlated with potential drug combinations that can be repurposed for targeting genes with poor prognosis in PDAC. Overall, these studies provide a better understanding of the metabolic vulnerabilities in PDAC and will lead to novel effective therapeutic strategies.

**Author summary:** Pancreatic ductal adenocarcinoma (PDAC) is the most common type of pancreatic cancer, with late diagnosis, early metastasis, insufficient therapy response, and very low survival rates. Due to these challenges associated with the diagnosis and treatment of PDAC, it has been a research area of interest. With the goal of understanding the metabolic reprogramming in proliferating PDAC cells, we reconstructed healthy and PDAC models by incorporating patient transcriptomic data into a genome-scale global human metabolic model. Comparing the metabolic flux space for the reactions in the two context-specific models, we identified significantly divergent pathways in PDAC. These results allowed us to further investigate growth-limiting genes in PDAC and identify potential drug combinations that can be repositioned for treatment of PDAC.

## Introduction

Pancreatic ductal adenocarcinoma (PDAC), with poor prognosis, resistance to radio- and chemotherapy, and a five-year survival rate of only 8.2% is the most prevalent form of pancreatic cancer and the third-leading cause of cancer-related morbidity in the USA[1]. Its poor prognosis can be attributed to its complicated and multifactorial nature, especially the lack of early diagnostic markers as well as its ability to quickly metastasize to surrounding organs[2–4]. Additionally, high rates of glycolysis and lactate secretion are observed in PDAC cells, fulfilling the biosynthetic demands for rapid tumor growth[1]. The combined action of regulatory T cells (Treg), myeloid-derived suppressor cells (MDSCs), and macrophages blocks theCD8^+^ T cell duties in tumor recognition and clearance and, ultimately, results in PDAC cells manifesting extensive immune suppression[2].

PDAC microenvironment is greatly dominated by the presence of dense fibroblast stromal cells. In addition to creating an acidic extracellular environment, the dense stroma surrounding the tumor reduces oxygen diffusion into pancreas cells, resulting in hypoxia. In response to the reduced oxygen uptake, the tumor cells undergo metabolic reprogramming to favor Warburg effect metabolism^12^, which involves increased rates of glycolysis. Because cancer cells are characterized by unregulated growth, much of the cellular metabolism is hijacked to maximize the potential to generate biomass. Since PDAC cells are forced to live within a particularly severe microenvironment characterized by relative hypovascularity, hypoxia, and nutrient deprivation, these must possess biochemical flexibility in order to adapt to austere conditions. Rewired glucose, amino acid, and lipid metabolism and metabolic crosstalk within the tumor microenvironment contribute to unlimited pancreatic tumor progression. The metabolic alterations of pancreatic cancer are mediated by multiple factors. These cells survive and thrive mainly in three ways: (1) Reprogramming intracellular energy metabolism of nutrients, including glucose, amino acids, and lipids; (2) Improving nutrient acquisition by scavenging and recycling; (3) Conducting metabolic crosstalk with other components within the microenvironment[5]. In addition, the metabolic reprogramming involved in pancreatic cancer resistance is also closely related to chemotherapy, radiotherapy and immunotherapy, and results in a poor prognosis. Thus, investigations of metabolism not only benefit the understanding of carcinogenesis and cancer progression but also provide new insights for treatments against pancreatic cancer. A better understanding of the metabolic dependencies required by PDAC to survive and thrive within a harsh metabolic milieu could reveal specific metabolic vulnerabilities.

Systemic chemotherapy is presently the most frequently adopted treatment strategy for PDAC. However, chemotherapy treatments often show limited success due to intrinsic and acquired chemoresistance[6, 7]. While many previous studies have predicted potential biomarkers for therapeutic purposes, including the *ribonucleotide reductase catalytic subunits M1/2 (RRM1/2)*, an enzyme catalyzing the reduction of ribonucleotides, or the *human equilibrative nucleoside transporter 1 (hENT1)*, a transmembrane protein, the treatment with drugs (i.e., gemcitabine and other combinatorial drugs) often failed [8–12]. The hypoxic microenvironment is also resistive to radiation dosage, reducing the efficacy of radiotherapy. In addition, the overexpression of key regulators of the DNA damage response (e.g., *RAD51* in PDAC) has been reported to contribute to the accelerated repair of DNA damage [128, 129]. Several genes have been reported to be frequently mutated in PDAC (*i.e., KRAS, CDKN2A, TP53*, and *SMAD4*)[13, 14] and, therefore, received increased attention as potential drug targets [15–19]. However, successful therapeutic strategies are yet to be developed [20–22]. The downstream events of metabolic reprogramming are considered as prominent hallmarks of PDAC[23]. Therefore, tackling this aggressive cancer through establishing a clear understanding of its metabolism has been a critical challenge to the scientific and medical communities. Since the underlying mechanism of these drug-resistive metabolic traits are only poorly understood, it warrants the use of novel computational techniques to understand the metabolic landscape of tumor progression and further compliment the going experimental efforts.

The increase in knowledge of macromolecular structures, availability of numerous biochemical database resources, advances in high-throughput genome sequencing, and increase in computational efficiency have accelerated the use of *in silico* methods for metabolic model development and analysis, biomarkers/therapeutic target discovery, and drug development[24–29]. These models provide a systems-level approach to studying the metabolism of tumor cells based on conservation of mass under pseudo-steady state condition. Since genome-scale metabolic models are capable of efficient mapping of the genotype to the phenotype [30–35], integrating multi-level omics data with these models enhances their predictive power and allows for a systems-level study of the metabolic reprogramming happening in living organisms under various genetic and environmental perturbations or diseases. Applications of the genome-scale metabolic modeling to cancer includes network comparison between healthy and cancerous cells, gene essentiality and robustness studies, integrative analysis of omics data, and identifying reporter pathways and reporter metabolites [36–40]. For example, Turanli *et. al* used metabolic modeling to pinpoint drugs that could effectively hinder growth of prostate cancer[37]. Similarly, Katzir, et. al mapped the reactions and pathways in breast cancer cells using a human metabolic model and various “omics” datasets[41]. Pancreatic cell and pancreatic cancer metabolism have been modeled before as a part of reconstructing draft models of several human cell types aimed at identification of anticancer drug through personalized genome-scale metabolic models [28, 42]. Although a pan-cancer analysis of the metabolic reconstructions of ~4000 tumors were attempted recently [43], the models generated were tasked with only finding the origin of the cancer-specific genes and reactions, and were not essentially curated and refined to achieve a high level of predictability. Kinetic modeling of the pancreatic tumor proliferation was also attempted, by modeling the glycolysis, glutaminolysis, tricarboxylic acid cycle, and the pentose phosphate pathway to find enzyme knockout or metabolic inhibitions suppressing the tumor growth [44]. While these studies have advanced our understanding of the metabolic landscape of pancreatic ductal adenocarcinoma or cancer in general, there is still necessity of a highly curated and predictive genome-scale metabolic model in order to have a system-level understanding of the metabolic changes.

To understand the PDAC-associated metabolic reprogramming that involves changes in the metabolic reaction fluxes and metabolite levels, genome-scale metabolic reconstructions of the healthy human pancreas and the PDAC cells encompassing the genes, metabolites, and reactions, were developed. This reconstruction process utilized patient transcriptomic dataset from the Cancer Genome Atlas (https://www.cancer.gov/tcga). The models were used to elucidate the altered metabolism of PDAC cells compared to the healthy pancreas. A concise schematic of the workflow in this study is presented in Figure 1. Upon incorporation of the transcriptomic data, the shifts in reaction flux spaces were observed across the metabolic network, notably in glycolysis, pentose phosphate pathway, TCA cycle, fatty acid biosynthesis, Arachidonic acid metabolism, carnitine metabolism, cholesterol biosynthesis, and ROS detoxification metabolism. Many of the observed metabolic shifts are in accordance with previously identified cancer hallmarks in omics-based studies. In addition, unique metabolic behavior was observed in mitochondrial and peroxisomal fatty acid beta oxidation, various parts of lipid biosynthesis and degradation, and ROS detoxification, which are discussed as potential for prognostic biomarkers. Significant downregulation in the peroxisomal fatty acid beta oxidation pathway reactions was observed in this study, which explains the shifts in cellular energy production and storage preference during pancreatic tumor proliferation. Furthermore, flux modulation in the carnitine shuttle system and the upregulation in the reactive oxygen species detoxification pathway reactions that was observed in this study indicate the unique strategies the PDAC cells adopt for survival. Potential drug repositioning and synergistic interaction between existing drugs that repressed the differentially expressed genes with poor prognosis in PDAC were identified. These findings manifest the predictive capabilities of genome-scale metabolic models at the reactome-level and can potentially direct new therapeutic approaches.

**Figure 1:**
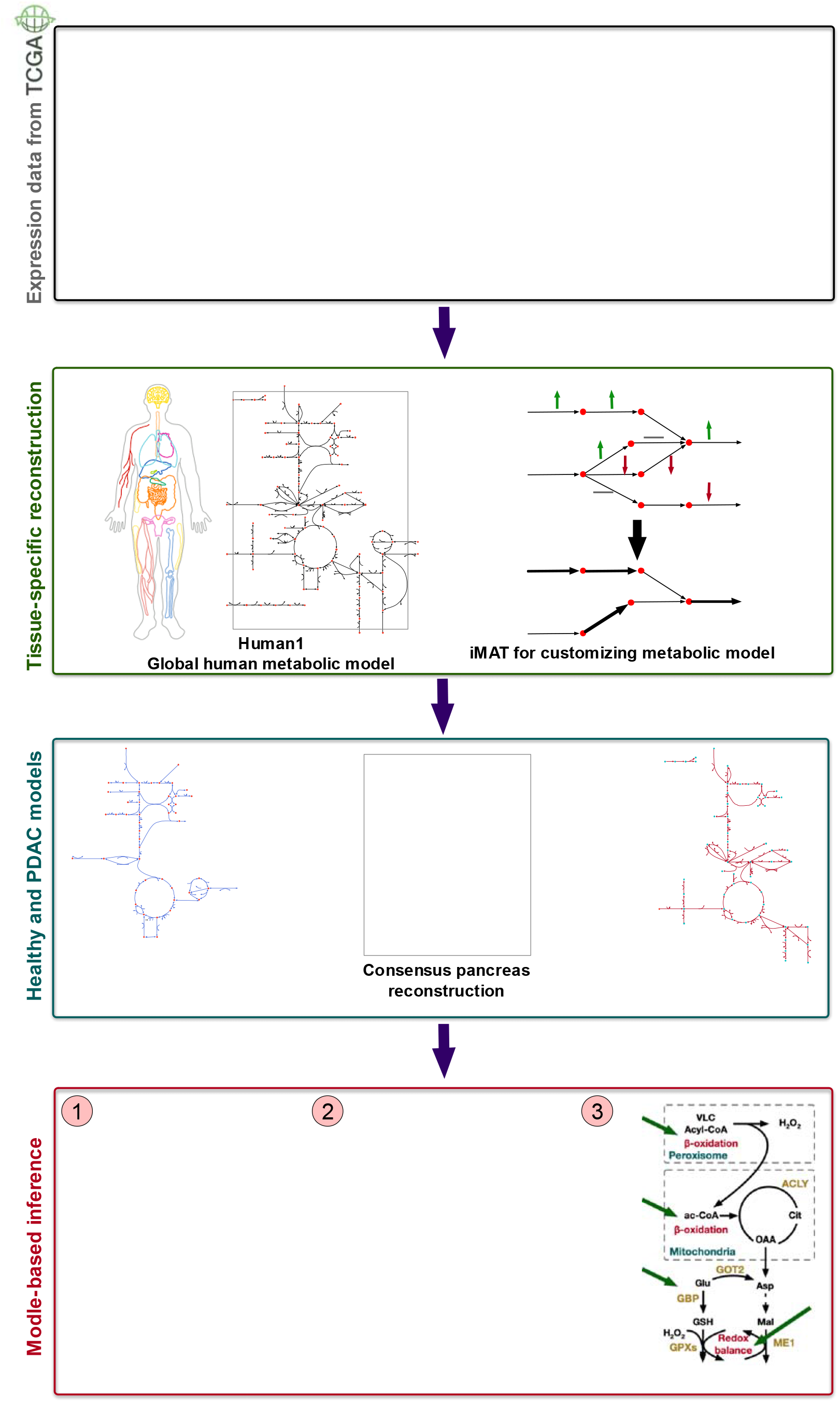
Schematic of the workflow for generating healthy pancreas and PDAC model and elucidating the metabolic divergence in PDAC.

## Results and Discussion

### Tissue-specific consensus pancreas metabolic reconstruction using transcriptomics data

A metabolic model describes reaction stoichiometry and directionality, gene-protein-reaction associations (GPRs), organelle-specific reaction localization, transporter/exchange reaction information, transcriptional/translational regulation, and biomass composition[45]. By defining the metabolic space, genome-scale metabolic models can assess allowable cellular phenotypes and explore the metabolic potential and restrictions under specific disease conditions[46]. The latest global human metabolic reconstruction, Human1[47], is an extensively curated, genome-scale model of human metabolism. It unified two previous and parallel model reconstruction lineages by the Systems Biology community, namely the Recon[48–50] and the Human Metabolic Reaction (HMR)[51, 52] series using an open-source version-controlled repository. In addition to curating the aggregated reconstruction, Human1 addressed issue with duplication, reaction reversibility, mass and energy conservation, imbalance, and constructed a new generic human biomass reaction based on various tissue and cell composition data sources. This standardized model allowed us to conveniently integrate omics data to develop a pancreas-specific metabolic reconstruction.

The transcriptomic data used to customize the global human model to a pancreatic reconstruction was obtained from the Cancer Genome Atlas (TCGA) database (https://www.cancer.gov/tcga). The Cancer Genome Atlas contains genomic, epigenomic, transcriptomic, and proteomic data on 33 cancer types in human, and is publicly available for the scientific research community. To obtain a representative set of transcriptomic data on both healthy and cancerous pancreas cells, 18 samples from the TCGA-PAAD project that contained quantified RNASeq transcriptomic data, were used. These samples contained Fragments Per Kilobase of transcript per Million mapped reads (FPKM) data of individuals from different ethnic backgrounds, ages, and sexes. Since the dataset accounted for a numerical expression value of every single of the 60483 genes across all the samples without any unique genes in the samples, the dataset was filtered for genes with no read count across samples. After that, 50392 genes remained, out of which 3628 metabolic genes overlapped with the genes in the Human1 metabolic reconstruction[47]. Differential gene expression analysis of the metabolic genes within the transcriptomic dataset from TCGA revealed 102 significantly differentially expressed genes, among which 53 showed significant upregulation and 49 showed repression in PDAC cells compare to healthy pancreatic cells (see details in Methods). Genes involved in glycolysis/gluconeogenesis, fatty acid and cholesterol biosynthesis, tRNA synthesis, Arachidonic acid metabolism, protein kinases, glutathione metabolism, RNA polymerase, DNA repair, mitochondrial beta oxidation, cytosolic carnitine metabolism, leukotriene and linoleate metabolism, and estrogen metabolism were consistently upregulated in all PDAC samples. On the other hand, genes related acylgylyceride metabolism, peroxisomal beta oxidation, mitochondrial and peroxisomal carnitine metabolism, several peroxidases, chondroitin, keratan, and heparan sulfate biosynthesis, glycerolipid metabolism, and different types of vitamin metabolism, including vitamins B12, D, and E, showed significant downregulation in PDAC. The complete results of differential gene expression analysis are presented in Supplementary information 1.

The preliminary pancreas metabolic reconstruction was obtained using the FPKM values for the 3628 metabolic genes in the TCGA dataset by iMAT[53] (details in the Methods section). It contained 3,628 genes, catalyzing 7,076 reactions, involving 4,415 metabolites located in 8 intracellular compartments (Cytosol, Mitochondria, Inner mitochondria, Golgi apparatus, Lysosome, Nucleus, Peroxisome, and Endoplasmic reticulum). The reactions are distributed across 133 different pathways, the largest of which include transport reactions, exchange/demand reactions, fatty acid oxidation, and peptide metabolism. Flux Variability Analysis[54] found that the 1444 reactions across 54 pathways could occur an unreasonably high rate not supported by thermodynamics, which are named unbounded reactions. The pathways contributing the largest number of unbounded reactions were transport, fatty acid oxidation, nucleotide metabolism, and drug metabolism. After the model had been refined by rectifying reaction imbalances and identifying and fixing infeasible cycles using Optfill[55] (see a complete list in Supplementary information 2), a thermodynamically feasible intermediate metabolic reconstruction of the pancreas encompassing all the reactions in both healthy and cancerous pancreas cells was obtained. This reconstruction was used as a baseline for generating the healthy and cancerous genome-scale pancreas metabolic model.

### Metabolic models of PDAC and healthy pancreas cell

The healthy pancreas and PDAC models were reconstructed from the consensus metabolic reconstruction of the pancreas. The Integrative Metabolic Analysis tool (iMAT)[53] was used to customize the model according to the gene expression values and corresponding ranking of the reactions (see methods section for details) in both healthy and PDAC cells. The healthy cell model contains 3,628 genes, catalyzing 6,384 reactions, across 129 pathways, involving 4,703 metabolites, while the PDAC cell model contains 3,628 genes, catalyzing 5,872 reactions, across 127 pathways, involving 4,381 metabolites. In both models, the pathways involving the largest number of internal reactions include fatty acid oxidation, cholesterol formation, peptide metabolism, and transport reactions. Supplementary information 3 and 4 contain the genome-scale metabolic model of the healthy and cancerous pancreas cells in Systems Biology markup Language level 3 version 1, respectively.

Figure 2 shows further details of the two models. While there are 5180 reactions overlap between the healthy and PDAC models, they have 1204 and 692 unique metabolic reactions, respectively (see Figure 2A and 2B). The unique reactions are distributed across divergent pathways in these two models (Figure 2C). The PDAC model distinctly shows better completeness of the Acyl-CoA hydrolysis, leukotriene metabolism, and starch and sucrose metabolism. On the other hand, many pathways have a more complete presence in the healthy cell model, including amino acid metabolism, structural carbohydrates (heparan and keratan sulfate) degradation, glycan metabolism, bile acid synthesis, and TCA cycle. While the more complete Acyl-CoA hydrolysis and sugar metabolism have been known to be associated with cancer cells, particularly interesting are the more complete leukotriene metabolism and lack of structural carbohydrate degradation pathways in the PDAC cell. It has been reported that the leukotrienes derived from membrane phospholipids play an important role in carcinogenesis [56, 57]. Furthermore, glycosaminoglycans (e.g., keratan sulfate, heparan sulfate, chondroitin sulfate) degradation in lysosomes are part of the normal homeostasis of glycoproteins. These molecules must be completely degraded to avoid undigested fragments building up and causing a variety of lysosomal storage diseases [58]. Lack of these degradation pathways in the PDAC indicate an increased accumulation of glycosaminoglycans in the tumor cell, which have previously been associated with cancer metastasis [59, 60].

**Figure 2:**
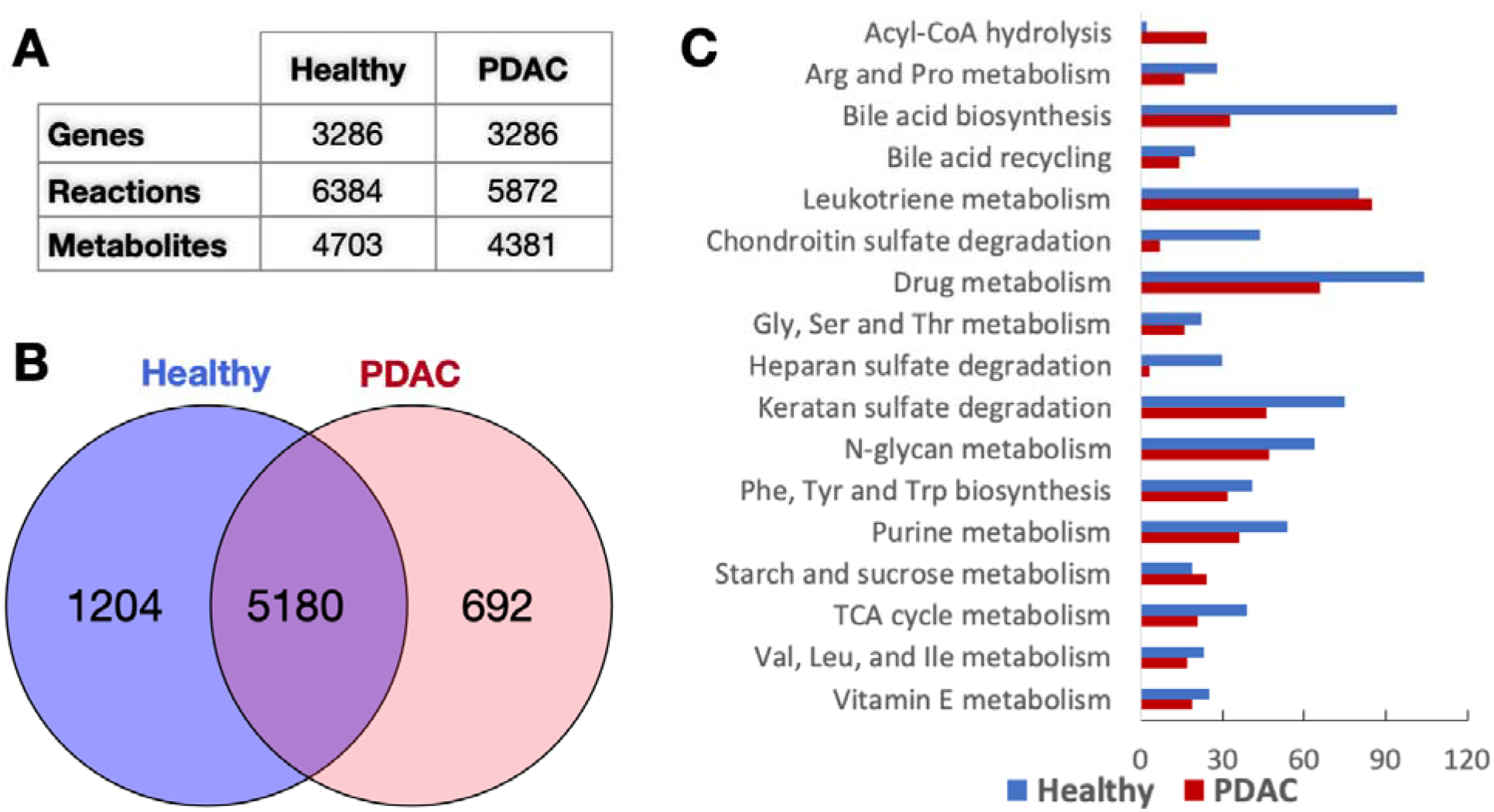
Model statistics for the healthy pancreas and the PDAC models. A) Numbers of Genes, Reactions, and Metabolites, B) overlap and uniqueness of metabolic reactions (Blue: Healthy, Red: PDAC), and C) Most divergent pathways between the two models.

### Unique metabolic traits in PDAC

The mathematically feasible flux ranges of the reactions in the healthy and PDAC models were assessed (see details in Methods sections) to explore the distinct shifts in PDAC cell metabolism. In Figure 3, the pathways with the biggest fraction of reaction fluxes significantly upregulated and downregulated are shown (a more detailed version is presented in Supplementary Information 5). While the observed metabolic shifts agree with the differential gene expression results discussed above, they also reveal some unique metabolic traits in PDAC. The model simulation results capture the most well-known metabolic hallmarks of pancreatic ductal adenocarcinoma. For example, the expansion of the flux space of the reactions in glycolytic pathways, bile acid biosynthesis, nucleotide metabolism, pentose phosphate pathway, and arachidonic acid metabolism is consistent with many studies[19, 23, 56] on pancreatic cancer in recent years.

**Figure 3:**
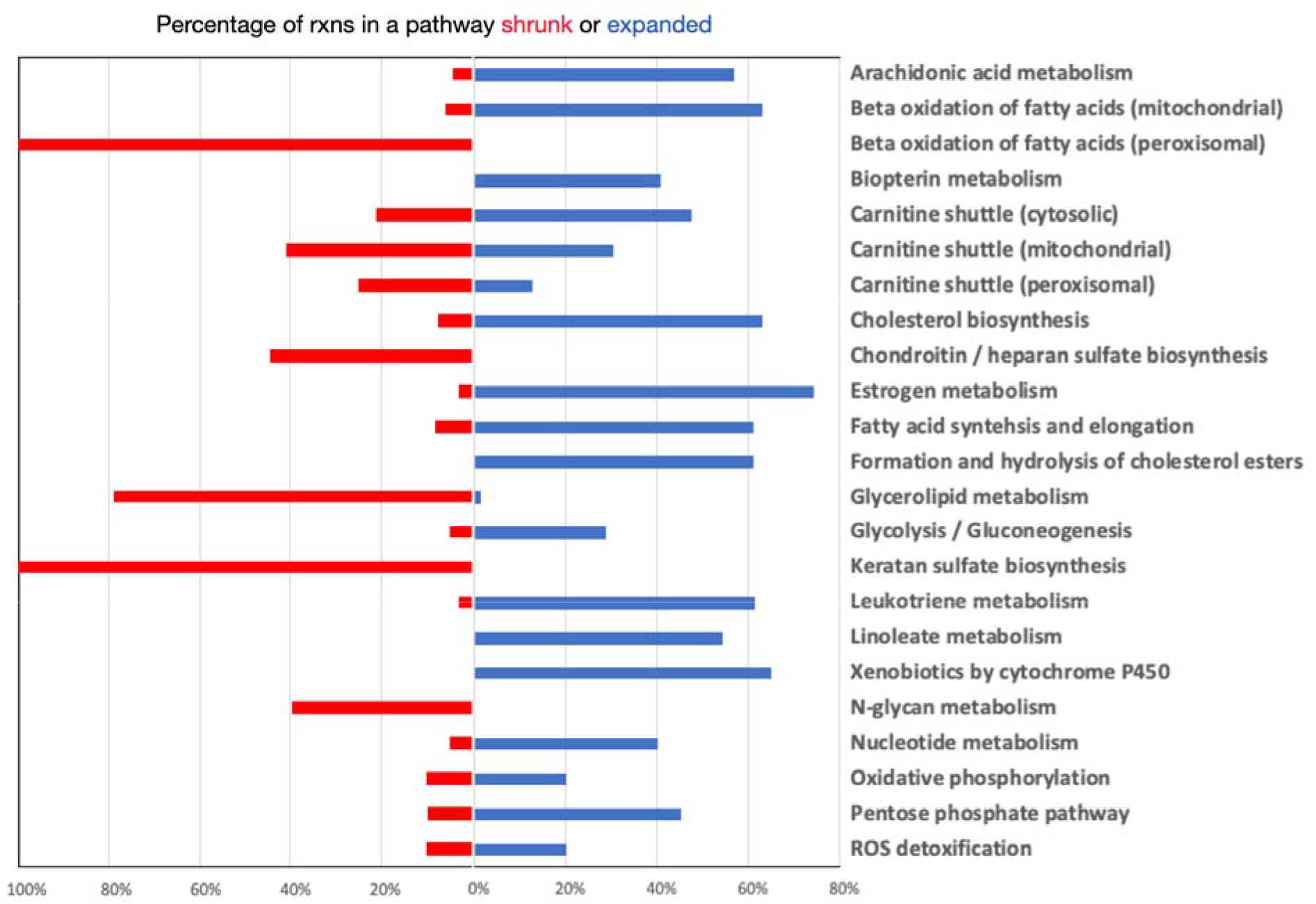
Significantly upregulated and downregulated pathways in PDAC cell metabolism. The bars (red: downregulated, blue: upregulated) represent the percentage of the total number of reactions in the respective pathway that changed their flux ranges.

These major metabolic reprogramming in pancreatic ductal adenocarcinoma arises from the well-known Warburg effect[61] due to constitutive activation of KRAS oncogene [62, 63]. KRAS activation in PDAC cells upregulates the uptake of glucose and enhance the glycolytic flux, including the production of lactate through lactate dehydrogenase (which demonstrates expanded flux ranges in PDAC) and channels carbon flux into the hexosamine biosynthetic pathway and pentose phosphate pathway. Both primary and metastatic PDAC tumors demonstrate increased glycolytic gene expression [64]. Notably, upregulation of pentose phosphate pathway and the downstream nucleotide biosynthesis pathway has been implicated in PDAC progression and therapy resistance [65–71]. Increased bile acid secretion has previously been identified in PDAC patients, which is indicative of tumor expansion into the bile duct[72] and may result in bile acid reflux into the pancreatic duct and acinar cells, from which PDAC is derived[73]. *NR1D1*, one of the two differentially expressed regulator genes, positively regulates bile acid synthesis[74], indicating a possible link between overexpression of that gene and PDAC carcinogenesis through increased bile acid synthesis. In addition, glutamine metabolism is vastly reprogrammed to balance the cellular redox homeostasis. Glutamine is sequentially converted to glutamate and aspartate in the mitochondria, which is shuttled into cytoplasm and eventually generates NADPH after a series of reactions to maintain redox homeostasis. The regeneration of NAD+ as an upstream substrate of NADH production is, therefore, an absolute requirement PDAC cell survival, particularly when mitochondrial demands escalate. Alterations in glucose and glutamine metabolism have also been linked with poor response to chemotherapy in PDAC [68, 71, 75].

Reactions in the arachidonic acid metabolism and leukotriene metabolism were observed to expand their flux space in PDAC. The two distinct branches of arachidonic acid metabolism, mainly driven by cyclooxygenase-2 (COX-2) and 5-lipoxygenase (5-LOX), were found to have significantly expanded their flux space in PDAC model. Several studies have reported that eicosanoid metabolism, especially arachidonic acid (AA) metabolizing enzymes including prostaglandins and leukotrienes (LT), play an important role in carcinogenesis [56, 57]. Specifically, the eicosanoids formed via COX-2 and 5-LOX metabolism directly contribute to pancreatic cancer cell proliferation in human[76]. Leukotrienes are also known to initiate inflammation and mount adaptive immune responses for host defense[77]. Prostaglandins have also been shown to regulate tumorigenesis in PDAC [78].

While the Warburg model explains these shifts to a great extent, especially in increased uptake of glucose and subsequent increased oxidative phosphorylation, recent studies have shown that the balance between glycolysis and oxidative phosphorylation may not always be in homeostatic. Rather, the metabolic reprogramming happening in PDAC is highly dynamic and dependent on the harsh tumor microenvironment [79]. Therefore, it is imperative to investigate other less suspected sources of unique metabolic traits of PDAC cells. Simulating the flux space of the PDAC cell model and comparing that with the healthy pancreas model allows us to examine the distinct changes metabolism in the reaction and pathway level. These observations are concisely presented in Figure 4.

**Figure 4:**
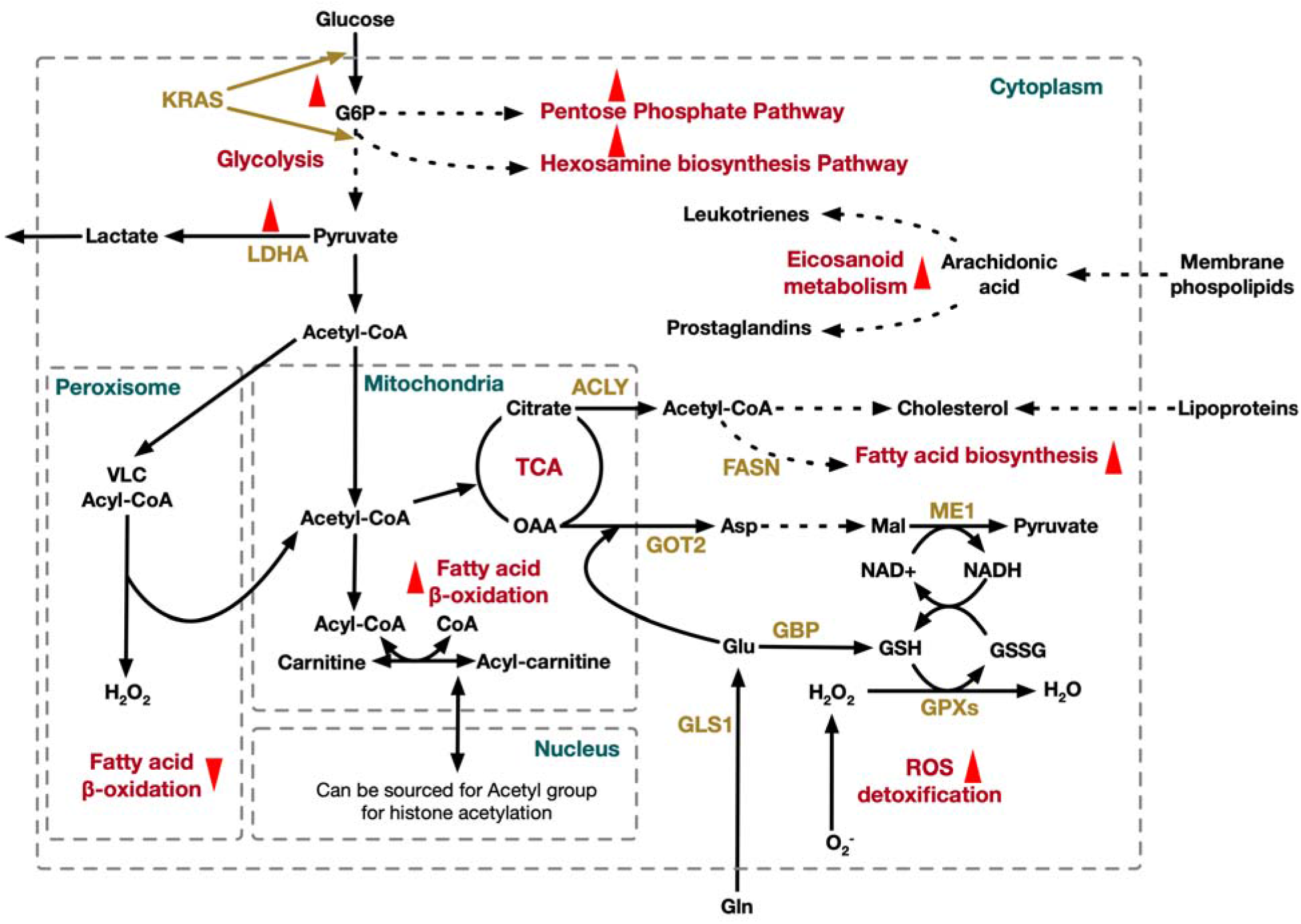
Distinct metabolic features of PDAC cell. ACLY: ATP-citrate lyase; Asp: aspartate; FASN: fatty acid synthase; Gln: glutamine; GLS1: glutaminase; Glu: glutamate; GOT: glutamic-oxaloacetic transaminase; GPX: glutathione peroxidase; GSH: glutathione reduced; GSSG: glutathione oxidized; LDHA: lactate dehydrogenase A; ME: malic enzyme; OAA: oxaloacetic acid; TCA: tricarboxylic acid; VLC: very long chain.

Increased abundance of acetyl-CoA and upregulated mitochondrial carnitine metabolism result in more carnitine and acyl-carnitine (mostly acetyl-carnitine) in the mitochondria. Carnitine can be transported to the cytosol and accumulated in biomass. Recent findings have suggested that carnitine shuttle could be considered as a gridlock to trigger the metabolic flexibility of cancer cells [80, 81]. Carnitine shuttle system is involved in the bidirectional transport of acyl moieties between cytosol to mitochondria, thus playing a fundamental role in tuning the switch between the glucose and fatty acid metabolism. This is crucial for the mitochondrial fatty acid beta-oxidation and maintaining normal mitochondrial function (balancing the conjugated and free CoA ratio) [82]. Higher burning of long-chain fatty acids produces increased energy for the cell to survive [83]. The available acetyl-CoA can be fed into the TCA cycle to produce more energy or acetyl moieties can be repurposed in the nucleus to recycle acetyl group for histone acetylation [84]. Thus, the carnitine shuttle system plays a significant role in tumor by supplying both energetic and biosynthetic demand for cancer cells[84].

While the mitochondrial beta oxidation pathway reactions primarily showed an expansion in flux space, all of the reactions in the peroxisomal beta oxidation pathway shrunk their flux space. This is an interesting feature of pancreatic ductal adenocarcinoma, since peroxisomal beta oxidation pathway was found to be upregulated in some cancer types [85] and downregulated in others [86–88]. The primary differences between fatty acid beta oxidation in mitochondria and peroxisome is the chain length at which fatty acids are synthesized and the associated product. Mitochondria catalyze the beta oxidation of the majority of the short to long-chain fatty acids, and primarily generate energy, while peroxisomes are involved in the beta oxidation of very-long-chain fatty acids and generate H_2_O_2_ in the process [89]. This means that while mitochondrial beta-oxidation is governed by the energy demands of the cells, peroxisomal beta-oxidation does not. Peroxisomal beta-oxidation is mostly involved in biosynthesis of very-long-chain fatty acids and do not produce energy, while the mitochondrial pathway is related to mostly catabolism and is coupled to ATP production [90]. Therefore, it is expected that the rapidly proliferating and energy-demanding tumor cells will favor the more energy-efficient mitochondrial pathways instead of the less required very-long-chain fatty acid-producing peroxisomal pathways. Furthermore, the reduction of the peroxide byproduct by downregulating the peroxisomal beta oxidation pathways reduces the oxidative stress, which helps the cancer cell to survive.

Lipid metabolism is essential for cancer progression since it provides the necessary building blocks for cell membrane formation and produces signaling molecules and substrates for the posttranslational modification of proteins. However, the role of fatty acids in pancreatic cancer is complicated and still not very well understood. In PDAC, we observe that reactions participating in *de novo* fatty acid biosynthesis, fatty acids elongation, and cholesterol biosynthesis pathways are upregulated, including citrate synthase, ATP citrate lyase, fatty acid synthase, and coenzyme A reductase. Overexpression of these lipogenic enzymes in PDAC have been reported in some previous studies as well [91–93]. Of note, increased fatty acid biosynthesis has been shown to impart poor chemotherapy responsiveness [93]. At the initial step of *de novo* lipid synthesis, ATP-citrate lyase (ACLY) converts citrate to acetyl-CoA, which is then channeled to cytoplasm. Acetyl-CoA and malonyl-CoA are coupled to acyl-carrier protein domain of fatty acid synthase (FASN) and the downstream genes to synthesize mono- and poly-unsaturated as well as saturated fatty acids [94]. Acetyl-CoA is also converted to cholesterol and cholesterol ester. This observation agrees with the elevated expression of HMG-CoA (3-hydroxy-3-methylglutaryl-Coenzym-A) reductase and LDLR (low density lipoprotein receptor) in a mouse model with PDAC [95]. In addition to higher intercellular lipid synthesis, uptake of extracellular lipids is also increased in PDAC. This indicates an increased demand of nutrients for rapid proliferation that the PDAC cells have to meet for *survival*.

Lactate dehydrogenase (LDHA) enzyme has shown a reversal of direction and increase in flux space in PDAC compared to healthy pancreas cell model, in the direction of lactate production. The overexpression of LDHA in pancreatic cancer and its ability to induce pancreatic cancer cell growth have been reported by Rong et al. in 2013 [96]. In addition, they showed that knocking down the LDHA in the pancreatic cancer cells significantly inhibited the cell growth revealing the oncogenic trait of LDHA and its association with poor prognosis [96]. LDHA overexpression and its association with the poor survival outcome have also been reported [97]. Although a complete mechanistic insight behind the causal effect of upregulation of LDHA could not be established yet, it potentially serves as an independent prognostic marker of PDAC.

## Potential Drug Repurposing

The uniqueness in gene expression and metabolic profile in PDAC cells allows for an extended search for potential drug-gene interactions. In addition to that, the ever-increasing challenges associated with the therapy-resistance of PDAC have necessitated the repurposing of old drugs. Leveraging the development in the various data-driven approaches, drug repurposing is becoming an efficient way of drug discovery which is cost effective. We identified 25 genes associated with poor prognosis in pancreatic cancer which had an overexpression in PDAC (see Supplementary information 6 for a complete list). In Figure 5, These genes are the shown to be associated with several drug currently in use in human, which are at different stages of the approval process. The edges connecting the drug to the genes indicated the evidence of repressive effects on the genes, according to DrugBank Pharmaco-transcriptomic database [98]. Several of these drugs have potential synergistic association between each other, as shown in Figure 5. These non-oncology drugs can potentially target not only known but also hitherto unknown vulnerabilities in pancreatic cancer. While many of the drugs are either approved (*e.g*., Ofloxacin, Ciprofloxacin) or at the investigational stage (*e.g*., Puromycin) for treating other diseases in the human body, some of these drugs (*e.g*., troglitazone) has been withdrawn from the market due to risk of severe liver failure that can be fatal [99, 100]. Nonetheless, they are still included in this association study, since newer studies have revealed anti-proliferative activities of the derivatives of this drug in other cancer types [101–104], which can result in an improved benefit-to-risk ratio for these drugs as well as suggest new drug combinations for reduced hepatotoxicity [105].

**Figure 5:**
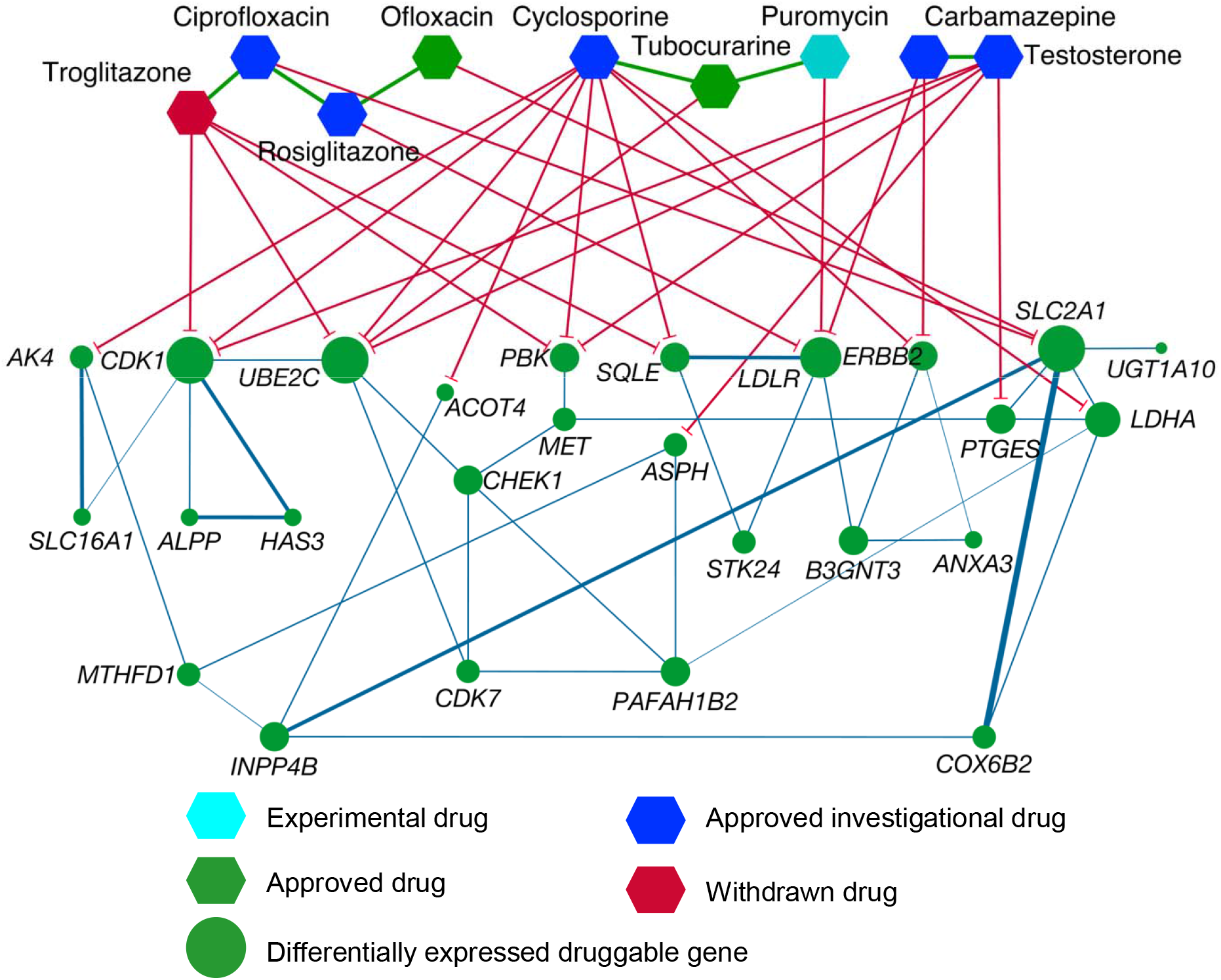
Potential drug interactions with upregulated genes in PDAC with poor prognosis. Edge thickness between the genes denote the correlation coefficient and the size of the nodes denote the magnitude of the gene expression fold change value in PDAC.

Since these drug-gene association are predicted in different tissue or disease systems and are a result of text mining through literature, we furthered our analysis of these associations by validating their effect on the fitness of the pancreatic cancer cell. To this end, we checked the inhibition effect on PDAC biomass when each of these genes are knocked out. The strongest growth inhibiting effect was observed when *SLC2A1* was knocked out, resulting in a no-growth phenotype during out model simulations. *SLC2A1* encodes major small sugar transporter across cellular membrane and between cellular organelles[106–110]. With its broad substrate specificity, *SLC2A1* can transport a wide range of aldoses including both pentoses and hexoses[110]. This is not only a rate limiting factor in sugar transport [107, 111], promoting aggressive tumor proliferation but also have been observed to be deregulated in pancreatic ductal adenocarcinoma [112]. Therefore, *SLC2A1* appears to be a high-confidence target for repositioning of the drugs repressing its expression, including fluoroquinolone-based antibiotics Ofloxacin and Ciprofloxacin. Other moderately growth-inhibiting genetic perturbations include monocarboxylate transporter *(SLC16A1)*, which is responsible for catalyzing the proton-linked transport of monocarboxylates such as L-lactate, pyruvate, and the ketone bodies [113]; Methylenetetrahydrofolate Dehydrogenase (*MTHFD1)*, which is closely coupled with nuclear *de novo* thymidylate biosynthesis [114]; and Cytochrome C Oxidase Subunit 6B2 (*COX6B2*), which accelerates oxidative phosphorylation, NAD+ generation, and cell proliferation [115].

To adapt to severe metabolic constraints, PDAC cells rely on specific metabolic reprogramming, thus offering innovative therapeutic strategies in the future. In this study, we attempted to identify a few poorly explored metabolic traits of PDAC cells, which can potentially complement the ongoing effort of finding novel therapeutic targets against pancreatic cancer. While many aspects of the pancreatic tumor progression have been studied with help of transcriptomics, proteomics, and metabolomics, this metabolic model-based study helps unravel the reactome layer of biochemical features that are associated with PDAC. While this systems-level metabolic analysis incorporates a relatively small sample size of clinical data, this allows us to assess the genome-scale changes in metabolism under tumor progression, and therefore can unravel previously unknown mechanistic insights into cancer cell proliferation as well as identify potential drug associations and synergistic drug combinations that can be repurposed. A better understanding of the metabolic dependencies needed to survive harsh conditions will uncover metabolic vulnerabilities and guide alternative therapeutic strategies.

## Methods

### Transcriptomic data processing

Transcriptomic data of 18 individuals (16 PDAC, 2 healthy normal) was obtained from the Cancer genome atlas (https://www.cancer.gov/tcga). The Fragments Per Kilobase of transcript per Million mapped reads were used as the input of differential gene expression analysis. The transcriptomic data included FPKM information for 60,483 genes for each of the samples. The FPKM values were filtered to exclude the genes with zero expression values throughout samples.

The DESeq algorithm in R software package “Bioconductor” was used for differential gene expression analysis [116]. DESeq employs negative binomial distribution and a shrinkage estimator for the distribution’s variance methods to test for differential expression [116]. Genes with a log2 (foldchange) value of 2 or higher were considered overexpressed and genes with a log2 (foldchange) value of −2 or lower were considered underexpressed, while satisfying an adjusted p-value of <0.05 [117]. Heatmap was generated using Morpheus (https://software.broadinstitute.org/morpheus) from the Broad Institute.

### Co-expression analysis with regulatory genes

Of the 490 differentially expressed genes, two over-expressed genes (*NR1D1* and *FOSL1*) were identified as regulatory genes using the Human Protein Atlas. A list of the genes regulated by each of these genes was obtained from RegNetwork[118]. The expression patterns of the two regulatory genes and their targets were examined to develop gene co-expression networks with the goal to identify highly co-expressed genes that could be considered regulators for genes expressed in PDAC. A threshold of >0.7 was used on Pearson’s correlation coefficient with a p-value of <0.05 for the development of the co-expression networks. Correlation clusters were developed grouping highly correlated genes to produce the co-expression networks. Network visualization was performed in Cytoscape[119] version 3.8.2 with manual repositioning. Gene expression data was visualized with varying node sizes, and correlation coefficients between genes were visualized with edge color and thickness.

### Preliminary pancreas metabolic reconstruction

A genome-scale metabolic model of a pancreatic cell describing reaction stoichiometry, directionality, and gene-protein-reaction (GPR) association was built by mapping these transcriptomic datasets to the latest global human metabolic model, Human1[47]. This global human model contains 13,417 reactions, 10,135 metabolites, and 3,628 genes, as of the github repository down in December 2020. This tissue-specific pancreas metabolic reconstruction was obtained from the Human1 model using the Integrative Metabolic Analysis Tool (iMAT)[53]. First, the reactions from the Human1 model were assigned artificial “expression values” (see Zur et al, 2010[53] for details) based on their associated gene and its corresponding expression values in the TCGA data. These expression values were then grouped into 3 categories: highly expressed, moderately expressed, and lowly expressed. Expression values greater than half a standard deviation above the mean were considered highly expressed and assigned a value of 1. Expression values less than half a standard deviation below the mean were considered lowly expressed and assigned a value of −1. Expression values that fell within a half a standard deviation of the mean were considered moderately expressed and assigned a value of 0. The expression for the Human1 biomass reaction was manually set to 1 so the biomass equation and all the other necessary reactions producing biomass precursors are included in the model. The iMAT algorithm then generated a model using the reaction expression information and reactions in the Human1 model.

### Flux Balance Analysis

Flux Balance Analysis (FBA)[120] was used to analyze the model performance during the different stages of refinement. The model was represented by a stoichiometric matrix, where the columns were representative of metabolites, and the rows representative of reactions. Constraints were imposed on the reactions given by upper and lower bounds for each based on nutrient availability and other conditions. FBA gives the flux value for each reaction in the model according to the following optimization formulation:

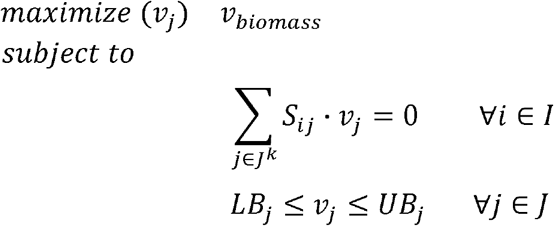

In this formulation, *I* is the set of metabolites and *J* is the set of reactions in the model. *S_ij_* is the stoichiometric coefficient matrix representing a model with *i* metabolites and *j* reactions, and *v_j_* is the flux value of each reaction. The objective function, *v_biomass_*, is representative of the growth rate of an individual cell. *LB_j_* and *UB_j_* are the minimum and maximum flux values allowed for each reaction.

### Model curation

The consensus model was curated through the classic design-build-test-refine cycle[121] to accurately reflect the metabolic capabilities of a pancreatic cell. Three reactions contained imbalances either in their stoichiometries or molecular formulas, and these imbalances were rectified. For reactions with imbalances caused by stoichiometric inaccuracies, changes were made to the stoichiometric coefficient matrix of the model. For reactions whose imbalances were due to incorrect molecular formulas, fixes were applied to the metabolic formula section of the model (see details in Supplementary information 2).

Thermodynamically infeasible cycles (TICs) are groups of reactions whose products, reactants, and directionality create a loop that allows unlimited flux to pass through each reaction, yielding no net consumption or production of metabolites. The presence of these cycles allows for many reactions in the model to occur at a very high rate even through the nutritional input to the model is negligible (or zero), which is unrealistic. These reactions are called unbounded reactions. It is important to eliminate these cycles to ensure the flux values for each reaction are thermodynamically feasible. Flux Variability Analysis (FVA) was performed on the model to identify mathematically possible flux ranges of the reactions in the model as well as identify the unbounded reactions. Unbounded reactions are characterized by flux distributions that hit the upper and/or lower bounds in FVA when all the metabolic uptake reactions are turned off. This initial analysis revealed 1444 unbounded reactions in the model, across multiple pathways including transport, fatty acid oxidation, nucleotide metabolism, and drug metabolism. The thermodynamically infeasible cycles comprising these unbounded reactions were identified using OptFill[55]. OptFill identifies TICs through iteratively identifying the smallest number of reactions with nonzero flux for which the sum of their fluxes is 0. All uptakes are turned off for OptFill so that all reactions carrying high flux are involved in a TIC. These cycles were eliminated by i) removing duplicate reactions from the model(s), ii) restricting reaction directionality if there is literature evidence of thermodynamic information, iii) removing erroneous reactions, and iv) using correct cofactors in reactions (for example NAD vs NADP) if that information is available. (complete details in Supplementary information 2). 932 reactions were modified in total. 609 reactions were turned off because they were duplicates of other reactions or lumped reactions. 23 reactions that were initially irreversible were made reversible if there was literature evidence indicating their reversibility. 286 reactions that were initially reversible were made irreversible in the forward direction, and 14 initially reversible directions were made irreversible in the backward direction. When turning reactions off to fix cycles, it was ensured that all essential reactions remained active in the model.

### Hypergeometric test for reaction enrichment analysis

Hypergeometric enrichment test was used to identify reaction pathways which are overrepresented in the set of reactions with altered flux space. The list of reactions with changing flux spaces obtained from running flux variability analysis was used to conduct a two-tailed hypergeometric test. This test was used to obtain the pathways showing significant representation in the list of altered reactions.

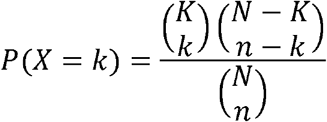

In this equation, *P(X=k)* is the probability that there are *k* reactions by chance with altered flux space in a given subsystem. *K* is the total number of reactions in a given subsystem, *N* is the total number of reactions in the model, and n is the total number of reactions in the model with altered flux space. The hypergeometric test was conducted for overrepresentation in each pathway in the model. For *P(X=k)* < 0.05, it is likely that the subsystem is over-represented due to a high number of altered reactions in the pathway rather than by chance. The p-values were then subjected to multiple-hypothesis correction using Benjamini-Hochberg method[122] using False Discovery Rate with α=0.05. From this, a list of pathways in the model most affected by PDAC was obtained.

### Drug interaction analysis

From the list of differentially expressed genes in the PDAC model, those associated with poor prognosis were identified using the Human Protein Atlas (http://www.proteinatlas.org). The differentially expressed genes associated with poor prognosis were then identified as potential therapeutic targets. For each of these genes, a list of drugs and their activation or repression effects were obtained from the DrugBank Pharmaco-transcriptomic database[98].

### Software and hardware resources

The General Algebraic Modeling System (GAMS)[123] version 24.7.4 was used to run FBA, FVA, and the OptFill algorithm on the model. GAMS was run on a high-performance cluster computing system at the Holland Computing Center of the University of Nebraska-Lincoln. The COBRA Toolbox[124, 125] version 3.0 in Matlab version 9.6.0.1174912 (R2019a) was used to run iMAT[53], identify essential reactions and reaction imbalances, and run FBA and FVA on the model.

## Author contributions

**Mohammad Mazharul Islam:** Data Curation, Formal Analysis, Investigation, Methodology, Resources, Software, Validation, Visualization, Writing – Original Draft Preparation

**Andrea Goertzen:** Formal Analysis, Methodology, Resources, Software, Writing – Original Draft Preparation

**Pankaj K. Singh:** Funding Acquisition, Writing – Review & Editing

**Rajib Saha:** Conceptualization, Funding Acquisition, Project Administration, Supervision, Writing – Review & Editing

## Data availability

All data generated or analyzed during this study are included in this published article and its supplementary information files.

## Supporting Information

Supplementary information 1: Differential gene expression analysis results.
Supplementary information 2: List of reactions removed, redirected, or balanced during model refinement.
Supplementary information 3: Genome-scale metabolic model of the healthy pancreas cell in Systems Biology markup Language level 3 version 1
Supplementary information 4: Genome-scale metabolic model of the PDAC cells in Systems Biology markup Language level 3 version 1
Supplementary information 5: Pathways with the biggest fraction of reaction fluxes significantly upregulated and downregulated
Supplementary information 6: Genes associated with poor prognosis in pancreatic cancer which had a significant differential expression in PDAC

