## Supplementary information 5 for "Exploring the metabolic landscape of pancreatic ductal adenocarcinoma cells using genome-scale metabolic modeling": Supplementary information 5.docx

Supplementary information 1

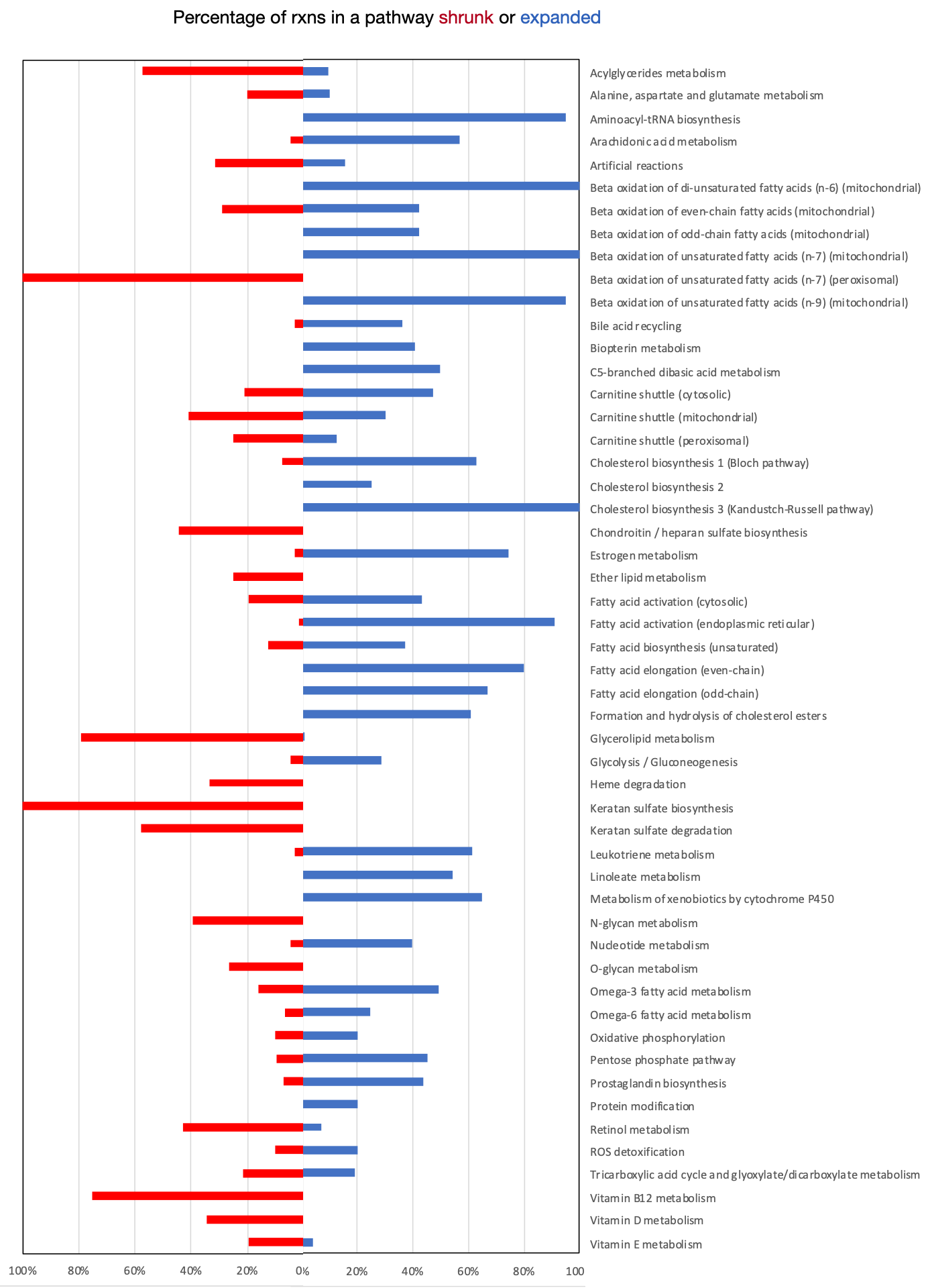

**Figure 1:** Significantly upregulated and downregulated pathways in PDAC cell metabolism. The bars (red: downregulated, blue: upregulated) represent the percentage of the total number of reactions in the respective pathway that changed their flux ranges.
